# How does cilium length affect beating?

**DOI:** 10.1101/474346

**Authors:** M. Bottier, KA. Thomas, SK. Dutcher, PV. Bayly

## Abstract

The effects of cilium length on the dynamics of cilia motion were investigated by high-speed video microscopy of uniciliate mutants of the swimming alga, *Chlamydomonas reinhardtii.* Cells with short cilia were obtained by deciliating cells via pH shock and allowing cilia to reassemble for limited times. The frequency of cilia beating was estimated from motion of the cell body and of the cilium. Key features of the ciliary waveform were quantified from polynomial curves fitted to the cilium in each image frame. Most notably, periodic beating did not emerge until the cilium reached a critical length between 2-4 µm. Surprisingly, in cells that exhibited periodic beating, the frequency of beating was similar for all lengths with only a slight decrease in frequency as length increased from 4 µm to the normal length of 10-12 µm. The waveform average curvature (rad/µm) was also conserved as the cilium grew. The mechanical metrics of ciliary propulsion: force, torque, and power all increased in proportion to length. Mechanical efficiency of beating appeared to be maximal at the normal wild-type length of 10-12 μm. These quantitative features of ciliary behavior illuminate the biophysics of cilia motion and, in future studies, may help distinguish competing hypotheses of the underlying mechanism of oscillation.

## Introduction

Motile cilia are highly conserved organelles that generate propulsive, oscillatory waveforms to propel cells or move fluids (1). The shape and characteristic frequency of the beating cilium are regulated by different components of its cytoskeletal structure: the 9+2 axoneme (2). The axoneme consists of a central pair of two singlet microtubules transiently connected by radial spokes to the surrounding nine doublet microtubules, which are in turn connected to each other by the nexin-dynein regulatory complex (nDRC). Minus-end directed dynein motor proteins anchored on one doublet exert forces on the neighboring doublet, which leads to relative sliding of the doublets and axonemal bending in the shape of propulsive waveforms (3, 4).

While the main structures of the axoneme have been identified by electron microscopy (5, 6) or cryo-electron tomography (cryo-ET) (7–11), the mechanisms that lead to a propulsive, oscillatory waveform remain incompletely understood. A number of distinct hypotheses have been proposed to explain the mechanism of waveform generation (12–17). Some of these hypotheses (13, 15–18) are formulated as mathematical models, which include length as a key physical parameter. Theoretical predictions of the effect of length on ciliary beat frequency and waveform are therefore possible. The current study focuses on experimental measurement of the effects of ciliary length on waveform during regrowth. These data can be compared to quantitative predictions from mathematical models and used to evaluate the underlying hypotheses. Questions addressed include: (i) Is a critical length required for periodic beating, and if so, what is it? (ii) How does length affect beat frequency? (iii) Does the shape of the waveform change as length increases, or does it simply scale? (iv) How does the mechanical output (force, torque and power) of the cilium change with length?

The unicellular alga*, Chlamydomonas reinhardtii*, which uses two cilia to swim toward a light source, is an excellent model system to study ciliary mechanics. Historically, these organelles were called flagella but the community henceforth agreed on using cilium and cilia for *Chlamydomonas*. Its ciliary beating is almost two-dimensional (2D) (19), which allows recording of the entire cilium within the focal plane of a standard optical microscope (20, 21). Wild-type *Chlamydomonas* assemble two cilia that are 10 to 12 µm long (22). Cilia of *Chlamydomonas* can experimentally be removed from the cell body using a variety of methods including mechanical shear or chemical stress (23, 24). The cells immediately begin to grow new cilia as the stress ends (25, 26). We chose to induce deciliation by subjecting the cells to an acid shock. This process allows use to record cilia at different length during the regrowth process. Cilia elongate up to its normal length in about 90 minutes (27). The *uni1* mutant strain assembles only one cilium, and the beating of its unique cilium creates a rotation of the cell body around an axis in a plane perpendicular to the plane of beating (28). The ciliary waveform of the *uni1* mutant slightly varies from the wild-type biciliate waveform as calculated from isolated axoneme (29, 30). Nevertheless, uniciliate cells show ciliary configurations similar to cilia of biciliated cells. (19, 28, 31)

Using a previously described method (20), we acquired and analyzed cilia waveforms. Briefly, ciliary motion is recorded with high-speed video-microscopy with a digital camera system and bright field optics. High-resolution, mathematical quantitative descriptions of the waveform of the cilia by a smooth surface of ciliary tangent angle can be extracted from the videos (Fig. 1).

**FIGURE 1:**
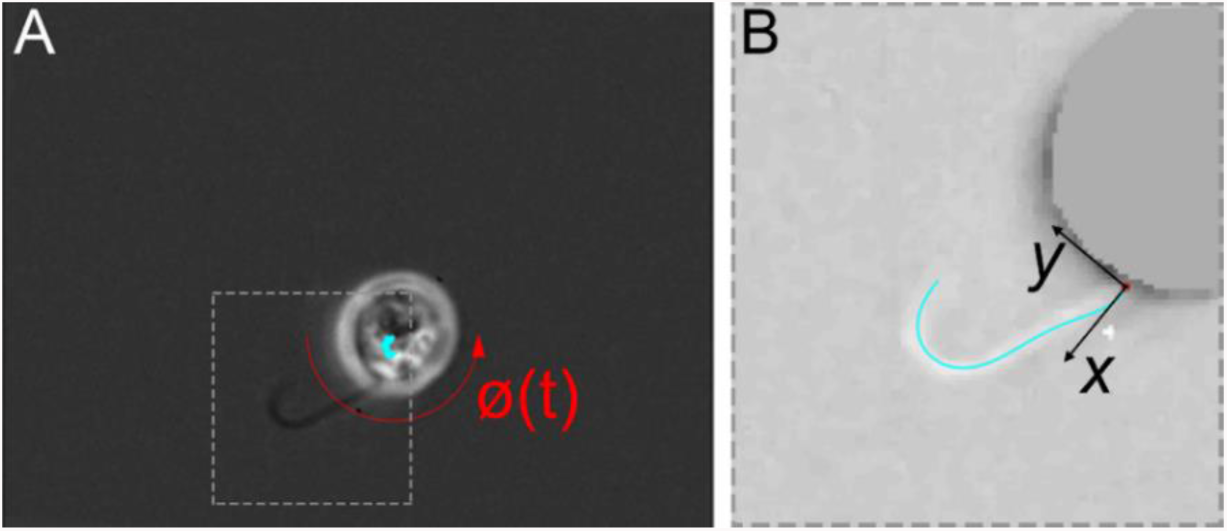
Example of analysis of body motion and cilium waveform. (A) The first video frame shows the translation of the center of the cell body (cyan curve) and cell body rotation ϕ(*t*) (red arrow) over the duration of the video. (B) Cropped area highlighting the cilium. The polynomial fit of the cilium is shown in cyan; its base is marked by a red circle. The Cartesian coordinate system based on the cilium proximal end is shown in black.

The evolution of the ciliary beat was quantified by a set of parameters that described the waveform as length increases. First, periodicity of the ciliary beating was determined from the normalized auto-covariance of the ciliary tangent angle. For periodically-beating cilia, beat frequency was measured from both body and cilia motion. In periodic cilia, a characteristic average waveform was defined and analyzed. Kinematic parameters (angle, curvature, and velocity) were extracted from the average waveform. Global force, torque, and power generated by the cilium were estimated from cilia motion (21) and compared to estimates from body motion. In addition, internal forces attributable to dynein motor protein activity were estimated from the waveform, using previous measurements of viscous resistive force coefficients, flexural modulus, and shear stiffness (32).

We find that periodic beating of cilia occurs only after a critical length is reached. Once beating begins, frequency is conserved, changing by only a few percent as length triples. The average curvature of the cilium is also strikingly consistent. Our analysis suggests that the intrinsic mechanics of the axoneme are stable during growth, and that increasing length alone can explain many qualitative changes in the waveform of the growing cilium.

## Materials and Methods

### Cell culture and deciliation

*Chlamydomonas reinhardtii* cells were grown as previously described (33). The uniciliate mutant strain *uni1-2* was generated from meiotic crosses, as described by Dutcher (34). Cells were grown on agar plate for 48 hours in Sager and Granick rich liquid medium supplemented with sodium acetate (35) at 25°C in constant light. Prior to recording, cells were suspended for 3 hours in a medium lacking nitrogen adapted from Medium I of Sager and Granick (35) to promote gametogenesis.

Short cilia were obtained by deciliation followed by regrowth for controlled duration. Deciliation was obtained by acid shock (24, 26). Medium (1 mL) containing cells was vortexed in an Eppendorf tube. Acetic acid (7 µL; 0.5 N) was added to the medium which was then vortexed for 45 seconds. Potassium hydroxide (3.5 µL; 0.5 N) was then used to buffer the solution. Finally, cells were vortexed (2g) for 3 minutes at 20°C and resuspended in rich liquid medium for ciliary regrowth. Cells were pipetted from the tube every 5 minutes during the 90 minutes of regrowth to be recorded under the microscope.

### Video-microscopy

All bright-field microscopy was carried out in a climate-controlled room maintained at 21°C. For each recording, 10 µL from medium containing cells were pipetted onto a slide, and a cover slip (18 × 18 mm) was placed for recording under a Zeiss Axiophot with a 100x Neofluar oil-immersion objective lens (Carl Zeiss AG, Oberkochen, Germany). Microscope settings were adjusted to provide the greatest contrast between the cilium and background of every single cell with a visible beating cilium. Videos were recording using a Phantom Miro eX2 camera and Phantom Camera Control Application 2.6 (Vision Research©, Inc, Wayne, NJ, USA). Videos were captured at 2000 frames per second with 320 × 240 resolution and an exposure time of 200 µs. Approximately 7000 frames were captured, with ~3500 frames before the trigger and ~3500 frames after the trigger. Roughly 1000 frames (0.5 sec) displaying typical beating were extracted and saved in uncompressed AVI format at 15 frames per second.

### Analysis Protocol

Videos were analyzed using a custom-made program written in Matlab™ R2016a (The Mathworks, Natick, MA, USA) modified from the version previously published (20). From each video, a sequence of 201 consecutive frames (0.1 sec) was stored as a 3D matrix of pixel intensity values. Each pixel had a spatial resolution of 169 × 169 nm and the temporal resolution between 2 consecutive time points was 0.5 ms. For slower moving cilia, in order to observe longer cycles of beating, movies were down-sampled up to 10 times to analyze a longer time interval.

Analysis of the ciliary waveform involved several steps. First, the motion of the cell body was characterized and a Cartesian frame (Χ, Υ) was defined based on the cilium proximal end (Fig. 1B). Then positions of points on the cilium were extracted from the video. After analysis of periodicity, in regularly-beating cells, an average characteristic beat was computed. Waveform kinematic parameters, global forces, and internal forces were then calculated as described in the following sections. Parameters and symbols used in the analysis are listed in Table 1.

**TABLE 1:** List of parameters and symbols.

| Symbol | Parameter | Unit |
| --- | --- | --- |
| $L$ | Length | $\mu\text{m}$ |
| $s$ | Axial coordinate along the cilium ($s = 0$ at base, $s = L$ at tip) | $\mu\text{m}$ |
| $\phi$ | Angle of body rotation | rad |
| $\Omega$ | Cell body average rotation rate | rps |
| $f_b$ | Frequency estimated from $\phi$ | Hz |
| $\theta$ | Tangent angle of cilium | rad |
| $a_c$ | Normalized auto-covariance of $\theta$ | |
| $\tau$ | Time lag in auto-covariance of $\theta$ | ms |
| $f_c$ | Ciliary beat frequency estimated from $\theta$ | Hz |
| $\kappa$ | Curvature, $\partial\theta/\partial s$ | rad-$\mu\text{m}$ |
| $F_{cx}, F_{cy}$ | Ciliary net force in $x$- and $y$-direction | pN |
| $P_c$ | Power generated by cilium | aW |
| $M_c$ | Torque applied by cilium, estimated from waveform | pN-$\mu\text{m}$ |
| $F_{bx}, F_{by}$ | Viscous force estimated from body motion | pN |
| $P_b$ | Power dissipated by body motion | aW |
| $M_b$ | Torque estimated from body motion | pN/$\mu\text{m}$ |
| $EI$ | Bending rigidity | pN-$\mu\text{m}^2$ |
| $k_\theta$ | Shear stiffness | pN/rad |
| $F_{dynein}$ | net dynein force (moment per unit length) | pN |
| $F_{bending}$ | force required to produce elastic bending of doublets | pN |
| $F_{viscous}$ | force required to overcome viscous drag | pN |
| $F_{shear}$ | force required to overcome resistance to shear | pN |
| $\overline{(\cdot)}$ | Average value of waveform parameter $(\cdot)$ | |
| $\widetilde{(\cdot)}$ | Amplitude (std. deviation) of waveform parameter $(\cdot)$ | |

### Analysis of cell body motion

Rigid-body motion of the cell was estimated as previously described (20). Briefly, each frame of the video was compared to rotated templates of the first image to define angles and displacements of the cell body (Fig. 1A). The rotation analyzed was always counter-clockwise; if the cell rotated clockwise, each video frame was flipped to reverse rotation direction. The cell body rotation rate Ω (revolutions per second, rps) and the ciliary beat frequency ƒ_b_ (Hz) were obtained from the time series of angular correction (ϕ). If the cell body did not rotate, Ω and ƒ_b_ were set to 0. If no peak was detected in the FFT of the rotation angle, ƒ_b_ was set to 0 as well.

### Mathematical description of the cilium position

Manual tracing followed by automated curve-fitting was used to provide a quantitative description of the cilium in space and time. As noted by others (36) fully automatic detection of *Chlamydomonas* cilia, especially shorter cilia, is difficult because of contrast variations; typically a bright “halo” surrounds the cell body. After correction of body motion, a rectangular region of interest (ROI) where the ciliary beating occurred was cropped by the user (Fig. 1A). The proximal end of the cilium (base) was defined manually by the user using a Matlab™ custom-made graphical interface.

A movie of the 199 frames displaying the ROI was saved in AVI format (uncompressed, 50 frames per second) for manual tracing of the position of the cilium. The user traced the position of the cilium on each frames of the video using ImageJ (37) brush tool (1-pixel width, cyan color) and an Intuos^®^ pen tablet (Wacom Technology Corporation, Vancouver, WA, Canada). The traced video was saved under AVI format (uncompressed, 15 frames per second)

The traced video was re-uploaded into Matlab™ and the cloud of traced points was stored as an array of Cartesian coordinates. The length of the cilium *L* was calculated and the points were fitted by a simple polynomial function θ(*s*) as described previously (20). The order of this polynomial fit was chosen as a function of the cilium length: for a cilium shorter than 3 µm, a second order polynomial function was used, a third order polynomial function was used for a cilium length between 3 and 6 µm and finally a fourth order polynomial for longer cilium. Each fit was performed using one point per pixel traced to calculate the fitting error. Varying the order of the polynomial with cilium length provided an approximately consistent ratio between the number of points used in the fit and the number of free parameters.

### Characteristic average beat

The auto-covariance *a*_*c*_ of the waveform angle *θ* was computed for each time point using the Matlab™ function *xcov*. The peak auto-covariance at non-zero lag 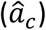 was used as a marker of periodicity and to find the period of ciliary beating (Fig. 2). Ciliary beat frequency *f*_*c*_ (Hz) is estimated from this period of beating. The theoretical value of 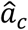 is 1.0 for perfect periodicity and each time point perfectly superimposed on each other; a peak of approximately 0.8 was typical of all periodic beats (Fig. 2A). If no peak was detected because none of signal were superimposing (Fig. 2B) or if the value of the peak was < 0.15, 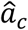 and *f*_*c*_ were both set to 0, and the cilium was considered non-periodic.

**FIGURE 2:**
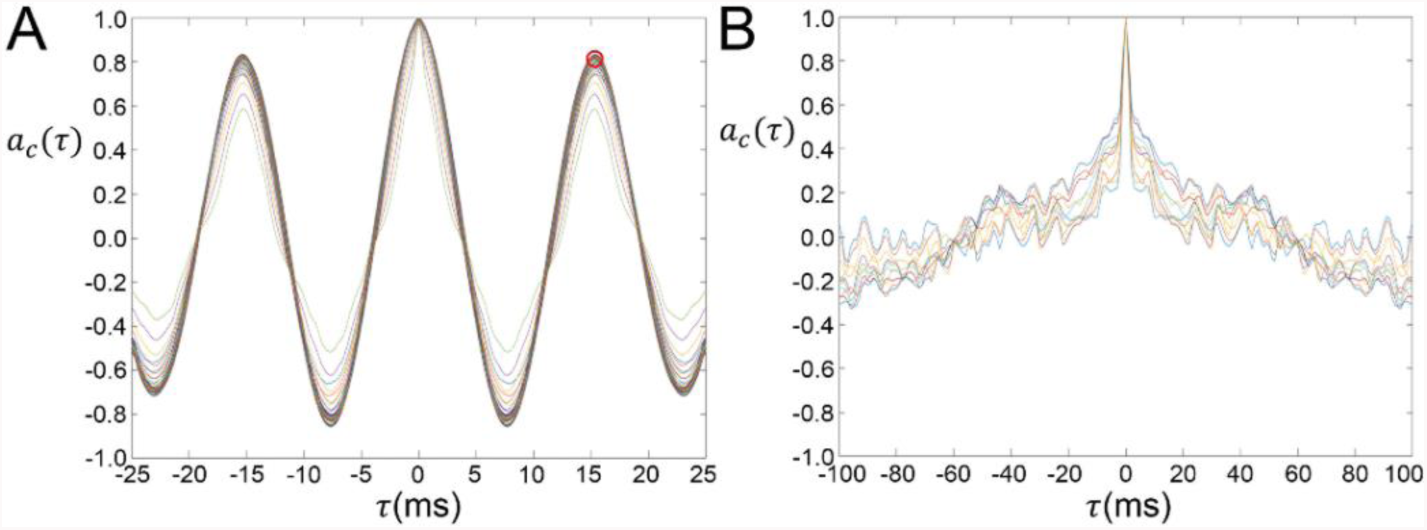
Quantification of periodicity of ciliary beating. Example plots of the normalized auto-covariance of cilia tangent angle, *a(τ)*, plotted vs. time lag, *τ*; the period of beating is defined by the first peak of magnitude greater than a specified threshold (0.15), at a non-zero lag 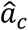 (red circle). (A) Example from cell with periodic beating, (B) Example from cell with non-periodic beating.

In periodically-beating cilia, values of the waveform angle *θ* from successive beats were averaged together to reconstruct a characteristic cycle of beating for which the parameters of waveforms could be calculated. We reconstruct a Cartesian coordinate (*x*, *y*) such as the proximal end of the cilium is (0,0) and the *x*-axis correspond to the direction of the forward swimming of a biciliated cell and the *y*-axis its normal (Fig. 1B).

### Waveform parameters and local forces

As described earlier (20, 29, 30), the surface *θ* (*s, t*) of the ciliary angle versus time and space can be used to extract key features illustrating the kinematics of the cilium. We calculated the average curvature 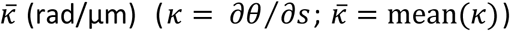, the bend amplitude 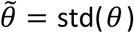 (rad), or the magnitude of the accumulated bend (which can be interpreted as a normalized average curvature) 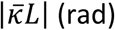.

The propulsive forces exerted by the cilium were estimated from the ciliary waveform. The normal and tangential components of the force per unit length applied by the cilium on the fluid (*f*_*N*_ and *f*_*T*_, pN/μm) are related by resistive force coefficients (*c*_*N*_ and *c*_*T*_) to the corresponding components of velocity (*f*_*N*_ = *c*_*N*_ *ν*_*N*_ and *f*_*T*_ = *c*_*T*_ *ν*_*T*_ (13, 38). Previously-obtained estimates of these coefficients (*c*_*N*_ = 0.0015 and *c*_*T*_ = 0.0007 pN-s/µm^2^) (21) are used in this study. The values of local resistive force can be transformed to Cartesian coordinates (*f*_*x*_, *f*_*y*_) and integrated to calculate the net forces of the cilium applied on the cell body in both *x*- and *y*-direction 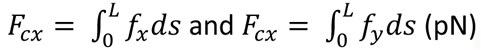. The power generated by the cilium 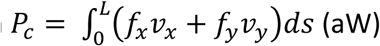 and the torque applied by the cilium to the cell body 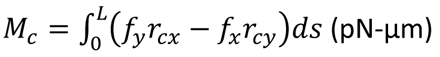 with *r*_*c*_ the relative positions of the point on the cilium with respect to the center of the cell body were calculated as well.

In the low Reynolds number regime, inertia may be neglected, so that the forces exerted by the cilium are balanced by the viscous drag of the body. Estimates of viscous forces and torque on the cell body (drag) and power dissipated by cell body motion were obtained from the formulas below for a prolate ellipsoid in Stokes flow (21, 39, 40).

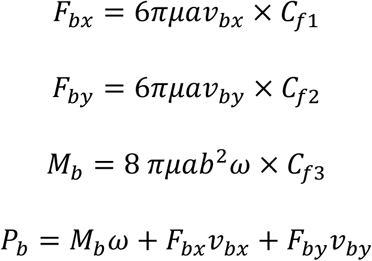

where *a* and *b* corresponded to the major and minor axis of the ellipsoidal cell body, *ν*_*b*_ the velocities of the body translation, *ω* the instantaneous angular velocity of the body rotation (*ω = dϕ/dt*) and the parameters *C*_*fi*_are functions of the eccentricity.

### Estimation of internal forces

Internal forces due to dynein motor protein activity are balanced by the elastic bending of the microtubule doublets and central pair, the viscous drag on the cilium from the surrounding fluid and the internal shear forces due to deformation of passive elements such as nexin links and radial spokes. Each process contributes a corresponding net moment (pN-µm) per unit length (µm). Since these quantities have units of force (pN-µm/µm = pN) they are represented by the symbols *F_*:

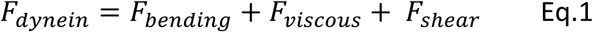

The internal force due to dynein activity can be estimated by calculating the terms on the right-hand side of (Eq.1) using approximate relationships and values from prior studies. Elastic bending moment per unit length, *F*_*bending*_, can be directly calculated from the local derivative of curvature of the cilium and the flexural rigidity *(EI* pN-μm^2^). The local viscous contribution to bending, *F*_*viscous*_, at any axial location on the cilium is calculated by integrating the viscous force per unit length from that location to the distal end of the cilium and taking the normal component, *F*_*N*_. Finally, elastic shear, *F*_*Shear*_, can be estimated from the product of the tangent angle of the cilium and shear stiffness *(K*_*θ*_, pN/rad).

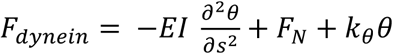

Bending rigidity *(EI)* and shear stiffness (*K*_*θ*_) were measured in wild-type cilia by Xu, *et al.* (32). As a first-order approximation for estimation of internal dynein force, all cilia were assumed to have the flexural rigidity and shear stiffness previously measured in normal length wild-type cilia, respectively *EI* = 840 pN-µm^2^ (mean ± std. dev. = 840 ± 280 pN-µm^2^) and *K*_*θ*_ = 40 pN/rad (mean ± std. dev. = 39.3 ± 6.0 pN/rad) (32, 41).

### Statistical analysis and comparison to predictions of mathematical models

For a given waveform parameter,(·), the average value, denoted by an overbar 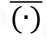, and the amplitude of its oscillations (RMS, or standard deviation), denoted by a tilde 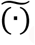, are estimated. All cells (deciliated or not) were considered as one group for statistical analyses. We created a classification of seven groups, based on ciliary length, with 2 µm sub-range intervals (Table 2). At least 50 beating cilia were recorded for each sub-range of length. Anova on ranks statistical tests were performed between the different sub-range of length using SigmaPlot (Systat Software, San Jose, CA). Significant differences reported in the manuscript have a p-value < 0.05.

**TABLE 2.** Number of movies and statistics for cilia length with different lengths.(n=388)

| Length range ($\mu\text{m}$) | Number of movies | Length mean and std. dev. ($\mu\text{m}$) |
| --- | --- | --- |
| [0, 2] | 51 | $1.5 \pm 0.3$ |
| [2, 4] | 57 | $2.9 \pm 0.6$ |
| [4, 6] | 53 | $5.2 \pm 0.5$ |
| [6, 8] | 59 | $7.2 \pm 0.5$ |
| [8, 10] | 63 | $9.0 \pm 0.5$ |
| [10, 12] | 55 | $11.0 \pm 0.6$ |
| [> 12] | 50 | $13.2 \pm 1.0$ |

In addition, a limited set of simulations were performed in order to help interpret experimental observations in the context of different hypotheses of the mechanism of ciliary oscillation. Four mathematical models, in the form of partial differential equations (PDEs) that describe the bending of thin elastic beams immersed in viscous fluid and driven by internal, active dynein force, were simulated. Modeling and simulation methods, references, parameter values, and results for each model are summarized in the Supporting Material.

## Results and Discussion

### Data overview

We recorded and analyzed 388 videos of beating cilia of uniciliate *Chlamydomonas* cells. Only cilia with apparent beating were recorded. 151 movies of beating cilia were recorded after 3 hours of gametogenesis. The length of these cilia varied from 6.4 to 16.3 µm. To record a wider range of ciliary length, 237 movies of beating cilia during the regrowth process were recorded following the deciliation protocol described above. In these cells ciliary length varied from 0.8 to 11.9 µm. Due to the bright “halo” around the cell body, cilia shorter than one micron were seldom possible to record with enough contrast for analysis. The regrowth rate of cilia (length vs time) was compatible with rates reported in the literature (27, 42, 43) (results not shown). Characteristic videos of each sub-range of length (Movies S1-8) are available in the Supporting Material.

### Cell body motion

We analyzed body motion in all 388 movies of ciliary beating. The cell body rotation rate Ω significantly increased with length (Fig. 3A). The beat frequency *f*_*b*_ extracted from the body motion was typically equal to zero or significantly reduced for cilia shorter than 2 µm. For cilia longer than 4 µm, *f*_*b*_ reached values around 60 Hz (59.9 ± 14.4 Hz) (Fig. 3B). Results summarized in Fig. 3B suggest that a key transition occurs around 4 µm and that cilia longer than 4 µm consistently exhibit periodic beating.

**FIGURE 3:**
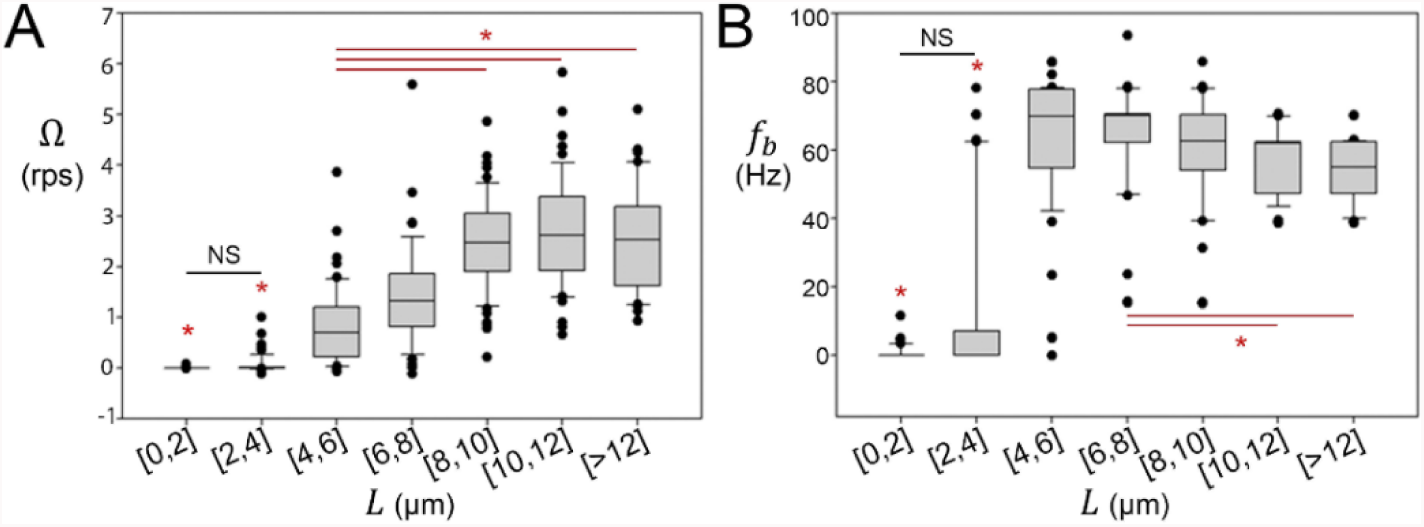
Analysis of body motion for each range of ciliary length. (A) Cell body average rotation rate Ω (revolutions per second, rps). (B) Beat frequency estimated from body motion *f*_*b*_ (Hz). (* significantly different (p<0.05), NS: Non-Significant. Statistical analysis by Anova on ranks).

### Periodicity of beating from analysis of cilia motion

We analyzed the periodicity of beating using the normalized auto-covariance of cilia angle, *θ*, from fitted waveforms (Fig. 2). Cilia shorter than 2 µm were never found to be periodic by this measure. Cilia between 2 and 4 µm length exhibited more variable periodicity. Cilia longer than 4 µm were usually periodic (no significant difference were found between the different length groups) (Fig. 4A). Those results suggest, again, that a change in ciliary behavior occurs around 4 µm. In total, 307/388 cilia analyzed were periodic. Non-periodic cilia included 51/51 cilia between 0 and 2 µm, 28/57 cilia between 2 and 4 µm and 2/53 cilia between 4 and 6 µm (Fig. 4B).

**FIGURE 4:**
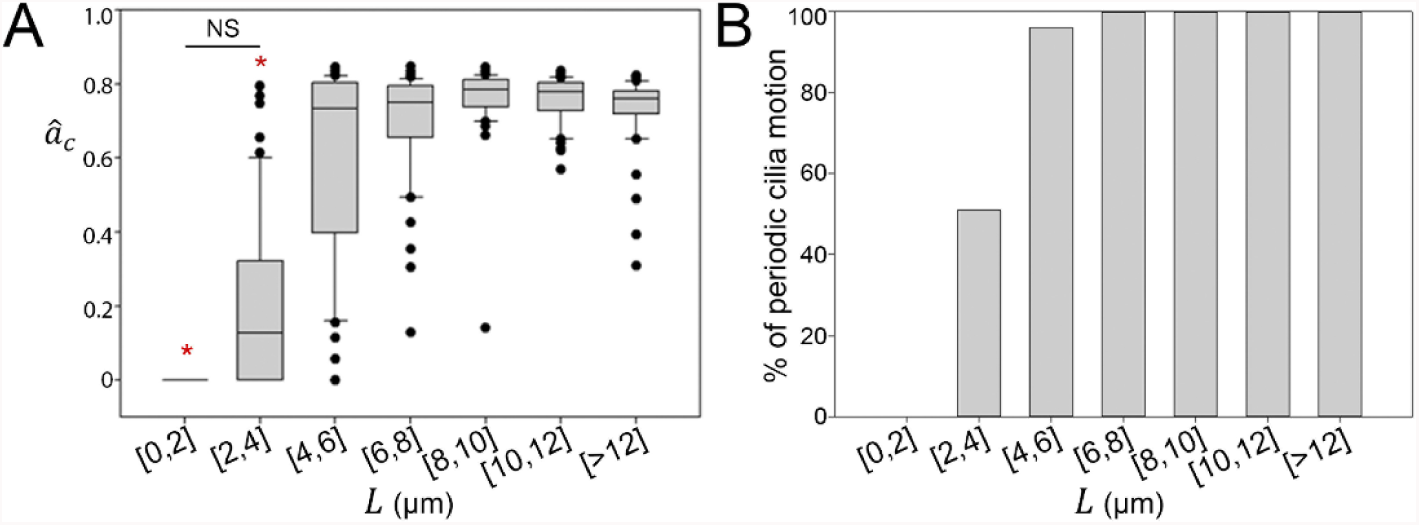
Analysis of periodicity of cilia motion for each range of ciliary length. (A) Value of the peak normalized auto-covariance 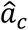 computed from waveform angle *θ*. Values <0.15 are considered to represent non-periodic behavior. (B) Percentage of periodic cilia motion.

### Waveform analysis during periodic beating

All waveform parameters reported are estimated for the characteristic beat, and are computed over the length of the cilium and the duration of the beat. Thus waveform parameters are only reported for periodic ciliary beating (*n*=307; cilia considered non-periodic are excluded).

Mathematical descriptions (polynomial coefficients of the average waveform) were obtained for 307 periodic-beating cilia. Fig. 5A shows a representative waveform for each group of length (respectively 3.3 µm, 5.1 µm, 7.1 µm, 9.1 µm, 11.0 µm and 13.1 µm). The specific waveforms in Fig. 5A were selected from each sub-range of length because the length *L*, the bend amplitude, 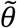, the average curvature 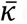 and the amplitude of oscillations in both *x*- and *y*-direction 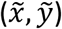 of each waveform were the closest to the average value within each respective sub-range. These plots represent the shape of the waveform at regular intervals of 1/10 of a cycle. The dimensional versions of the waveform (unscaled, Fig. 5A) illustrate the consistency in physical curvature in cilia of different lengths. The scaled waveforms (Fig. 5B) highlight qualitative differences in shape, particularly for cilia shorter than 4 µm.

**FIGURE 5:**
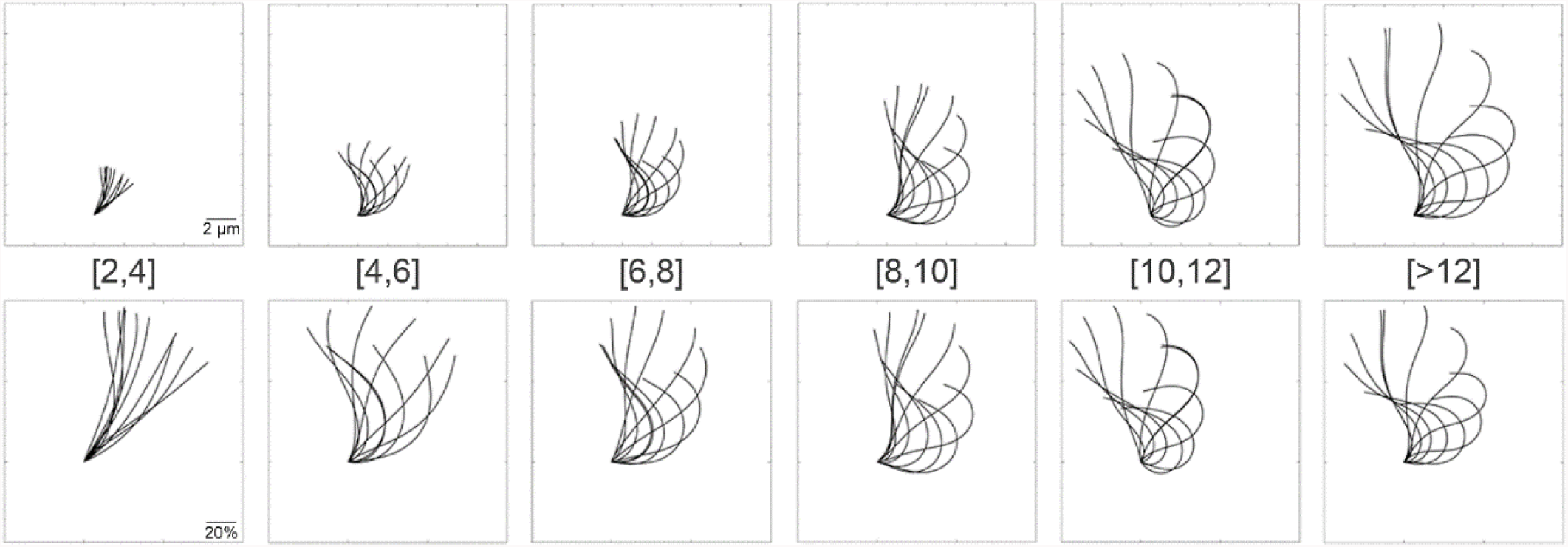
Representative ciliary waveforms for each length range. (Top row) Waveforms with dimensions in μm, reproduced to scale. (Bottom row) Dimensionless (scaled) waveforms.

### Waveform quantitative parameters

#### Beat frequency

The inverse of the time lag corresponding to the peak auto-covariance in cilia angle was used to quantify the ciliary beat frequency, **f**_*c*_, from waveform data. For longer cilia **f**_*c*_ was consistently near 60 Hz (mean ± std. dev. for all movies = 60.6 ± 13.6 Hz), exhibiting a slight decrease as length increased. Shorter cilia exhibit higher variability (Fig. 6A). This is consistent with prior estimates; the beat frequency in wild-type *Chlamydomonas* cilia has previously been reported to be 69.6 ± 8.7 Hz (20), 62.0 ± 2.7 Hz (44) or 63 ± 6 Hz (30). The beat frequencies estimated from cilia angle are also consistent with the frequencies independently estimated from the cell body motion (Fig. S1 in the Supporting Material). For cilia longer than 4 µm the ratio of frequency estimated from body motion to frequency of cilia motion *f*_*b*_/*f*_*c*_ is typically very close to 1.0 (mean ± std. dev 0.98 ± 0.15), as expected. Interestingly in short (2-4 µm) cilia the ratio was significantly different from 1 (Fig. S1B); apparently, even if the short cilium beats periodically, the frequency of beating is not reliably transmitted to the body motion.

**FIGURE 6:**
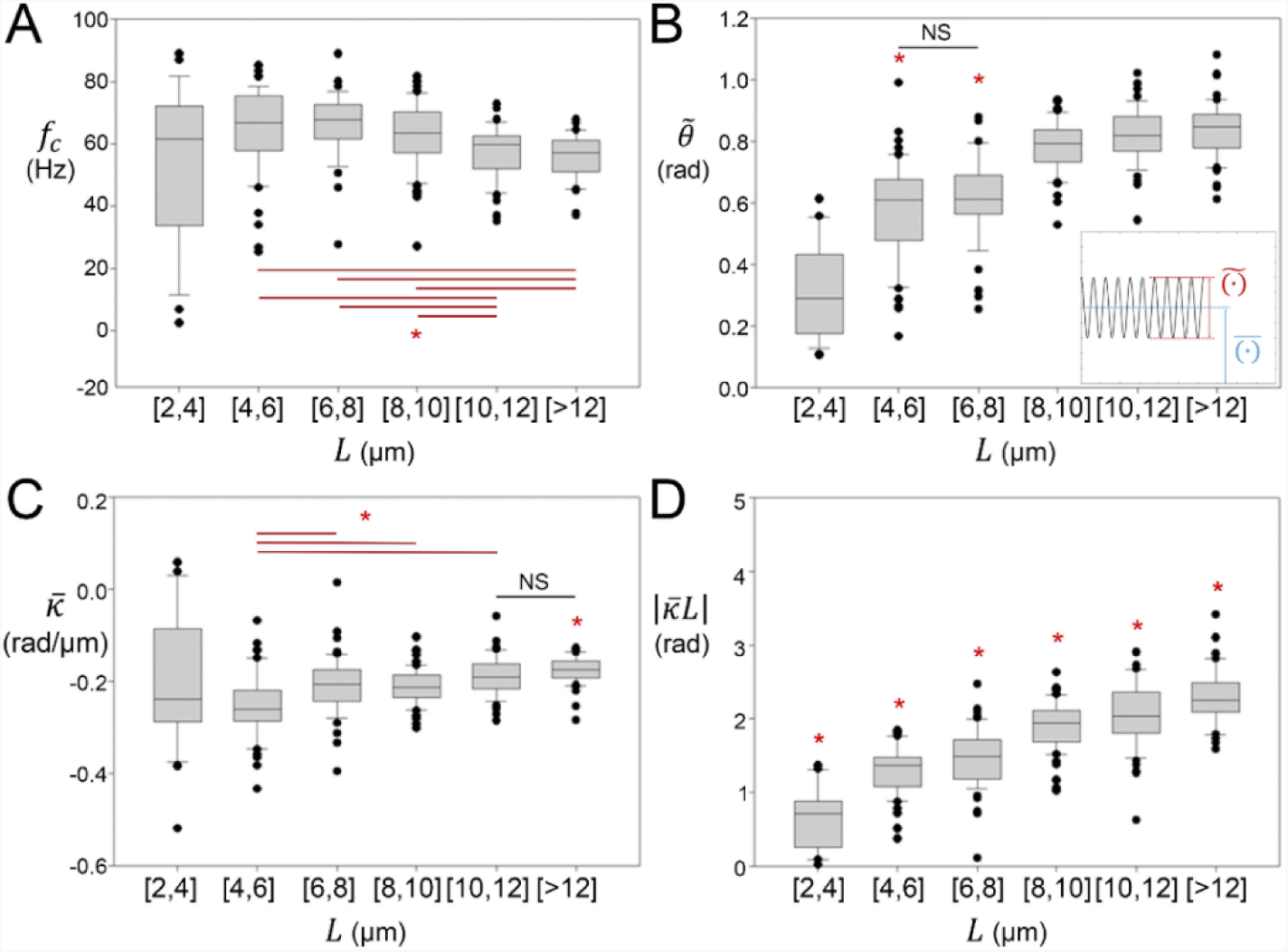
Parameters estimated from the tangent angle of the waveform, for periodically-beating cilia in each range of ciliary length. (A) Ciliary beat frequency *f*_*c*_ (Hz) estimated from tangent angle. (B) Bend amplitude 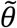 (rad). (Inset illustrates the mean value (blue, overbar) 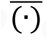 and amplitude (red, tilde) 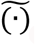 of a periodic parameter). (C) Average curvature 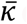(rad/μm). (D) Magnitude of accumulated bend (curvature normalized by length) 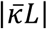 (rad).

#### Waveform shape

The bend amplitude 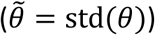 increased with cilium length (Fig. 6B). The average curvature 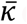(averaged over the length of the cilium and the full cycle of beating) was approximately conserved even for shorter cilia, at a value near −0.2 rad/µm (mean ± std. dev for all movies was −0.21 ± 0.07 rad/µm) (Fig. 6C). These values are consistent with previously-reported curvature values (− 0.17 ± 0.005 rad/µm for cilia 12.8 ± 1.5 µm long) (29, 30). The magnitude of the accumulated bend 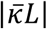 can be interpreted as a dimensionless curvature (curvature normalized by length). Because average curvature was fairly consistent at all lengths, 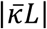 increased significantly with increasing cilia length (Fig. 6D). These observations are consistent with analogous observations of curvature and accumulated bend during maturation of cilia in human airway epithelial cells (45).

### Force, torque and power generated by the cilium

At low Reynolds numbers, the normal and tangential components of the force applied by the cilium to the fluid are related by resistive coefficients (*C*_*N*_ and *C*_*T*_) to the corresponding components of velocity (21), as described above. Integrating the normal and tangential components of the force over the cilium, we calculated the net force in *x*- and *y*-directions, respectively *F*_*cx*_ and *F*_*cy*_. The mean value and amplitude of force oscillations in both x- and y-directions increased with length (Fig. S2). More importantly for uniaxial cilium (since the cell rotates with little translation), the torque applied by the cilium to the cell body *M*_*c*_ and the power generated by the cilium *P*_*c*_ were also calculated. Both the mean value of torque 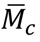 and the amplitude of torque oscillations 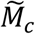 increased in proportion to length (Fig. 7A & B). The same trend was observed for the average power 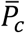 and the amplitude of oscillations of instantaneous power 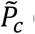(Fig. 7C & D).

**FIGURE 7:**
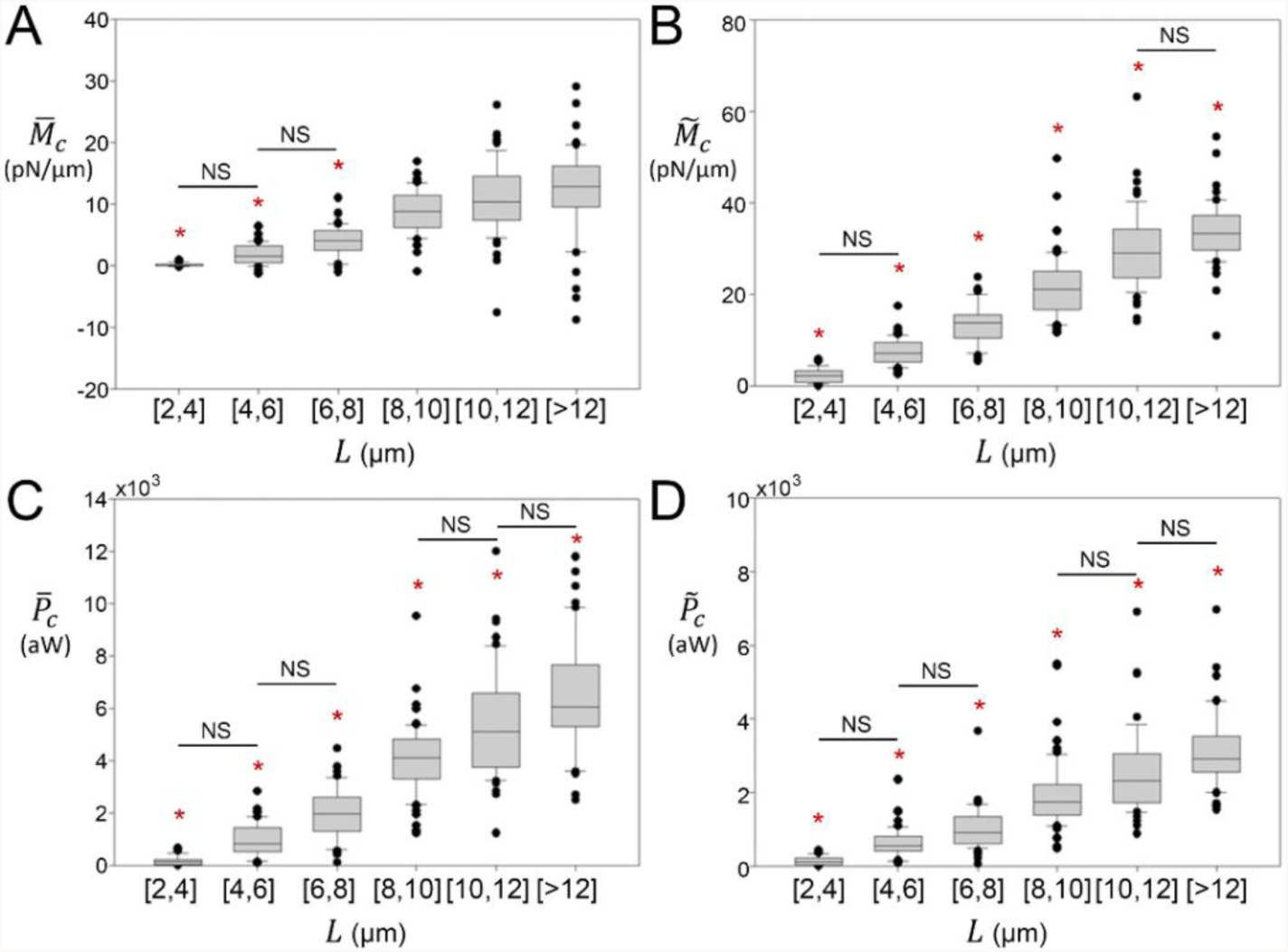
Global measures of power and torque developed by the cilium for each length range. (A) Average torque applied by the cilium about the center of the cell body 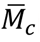 (pN- (B) Amplitude of variation in the instantaneous torque applied by the cilium to the cell body 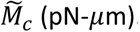. (C) Average power 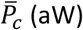 generated by the cilium. (D) Amplitude of variation in instantaneous power generated by the cilium 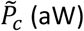.

Torque and power estimates derived from cilia motion and body motion are compared (Fig. 8). The torque generated by the cilium, *M*_*c*_, should balance the drag torque estimated from cell body rotation, *M*_*b*_. The torque estimates in Fig. 8A & B are approximately consistent with this principle, falling generally along the line of identity, although the lines of best fit suggest estimates of *M*_*b*_ are slightly (10-20%) higher than *M*_*c*_. This inequality may be due to small errors in the assumed resistive force coefficients, which were estimated in a previous study (21), or to the effects of outliers. The ratio 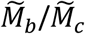 was around 1 in average (mean ± std. dev for all movies was 1.09 ± 0.67 and median was 1.01). In contrast, power dissipated by body motion, *P*_*b*_, is systematically much less than the power generated by the cilium, *P*_*c*_, reflecting the propulsive efficiency of this system (Fig. 8C & D). Notably, the average power dissipated by body motion *P*_*b*_ was around a third of the average power generated by the cilium *P*_*c*_. The ratio of the power amplitudes 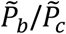 was close to 0.5 for all ranges of length (mean ± std. dev for all movies was 0.47 ± 0.29 and median was 0.42), although some differences were observed between the different length ranges. It is noteworthy that the highest ratio 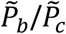 is observed for the (10-12 µm) cilia, which is the normal length range for *Chlamydomonas* (22).

**FIGURE 8:**
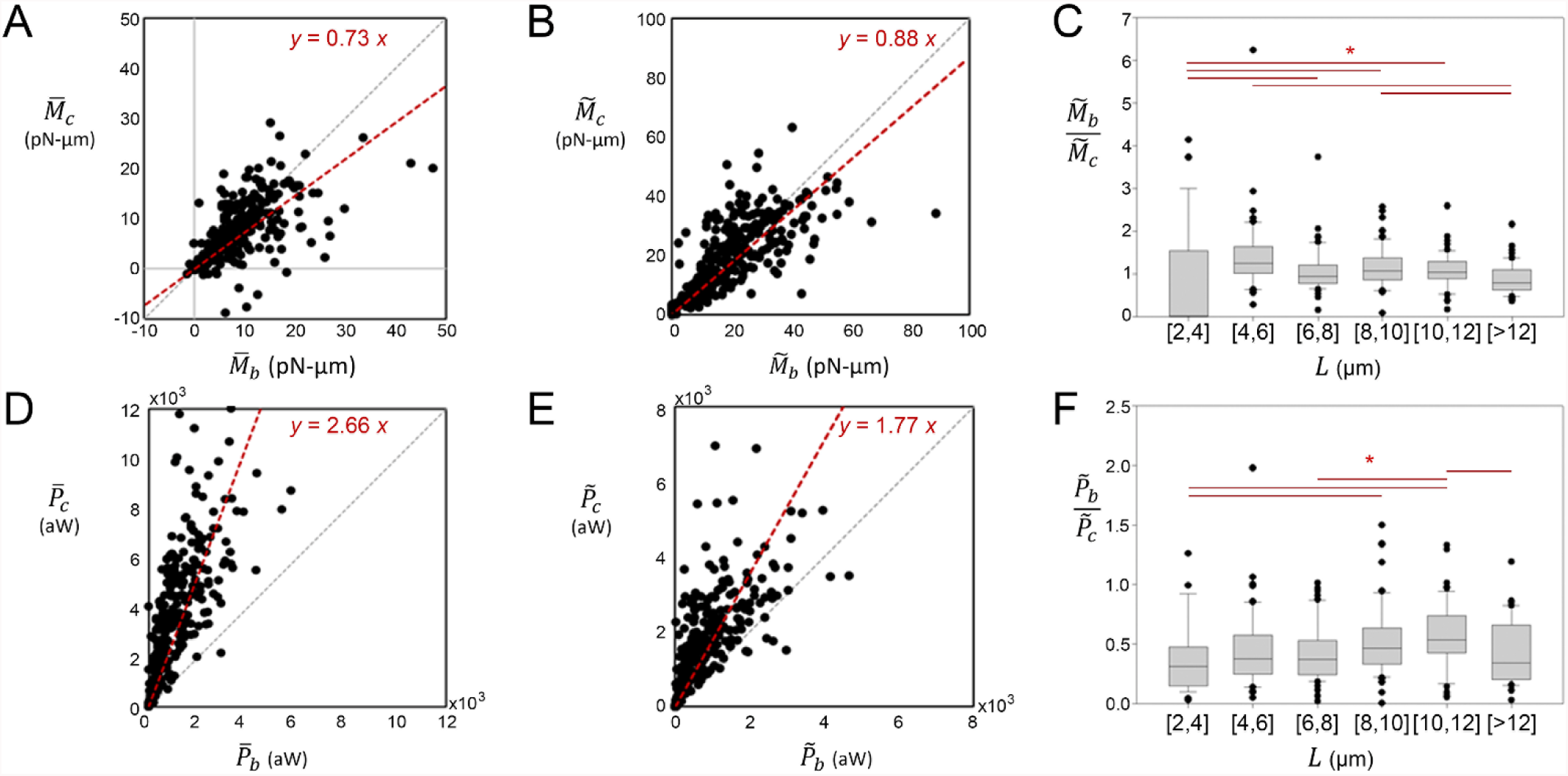
Comparison between torque and power estimates calculated from cell body motion and cilia waveform. (A). Average torque estimated from body motion 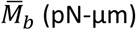 vs average torque estimated from waveform 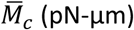. (B) Amplitude of variation in instantaneous torque estimated from cell body motion 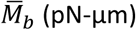 vs amplitude of torque estimated from waveform 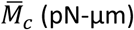. (C). Ratio 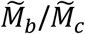 for each range of ciliary length. (D) Average power dissipated by cell body 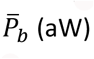 vs average power generated by the cilium 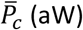. (E). Amplitude of variation in instantaneous power dissipated by cell body 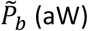 vs amplitude of power generated by the cilium 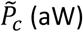 (grey dashed lines are the lines of identity, *y* = *x*; the red lines depict least-squares fits). (F) Ratio 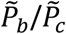 for each range of ciliary length.

### Internal forces

Internal forces within the cilium generated by dynein molecular motors were estimated from the contributions of the three separate opposing forces, which are viscous drag, elastic shear and elastic bending. While both drag and shear forces increase with cilium length, these components are much smaller than the force required to bend the doublets (Fig. 9). Thus, the net dynein force appears to be largely dedicated to overcoming the resistance to local elastic bending. For growing cilia of different length, the amplitude of oscillatory dynein force 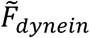(over the entire cilium during the full cycle of beating) was roughly conserved around 200 pN (mean ± std. dev 197.7 ± 52.9 pN). (We note that this observation relies on the assumption that the physical parameters: resistive force coefficients, shear stiffness and flexural rigidity, do not change as the cilium regrows).

**FIGURE 9:**
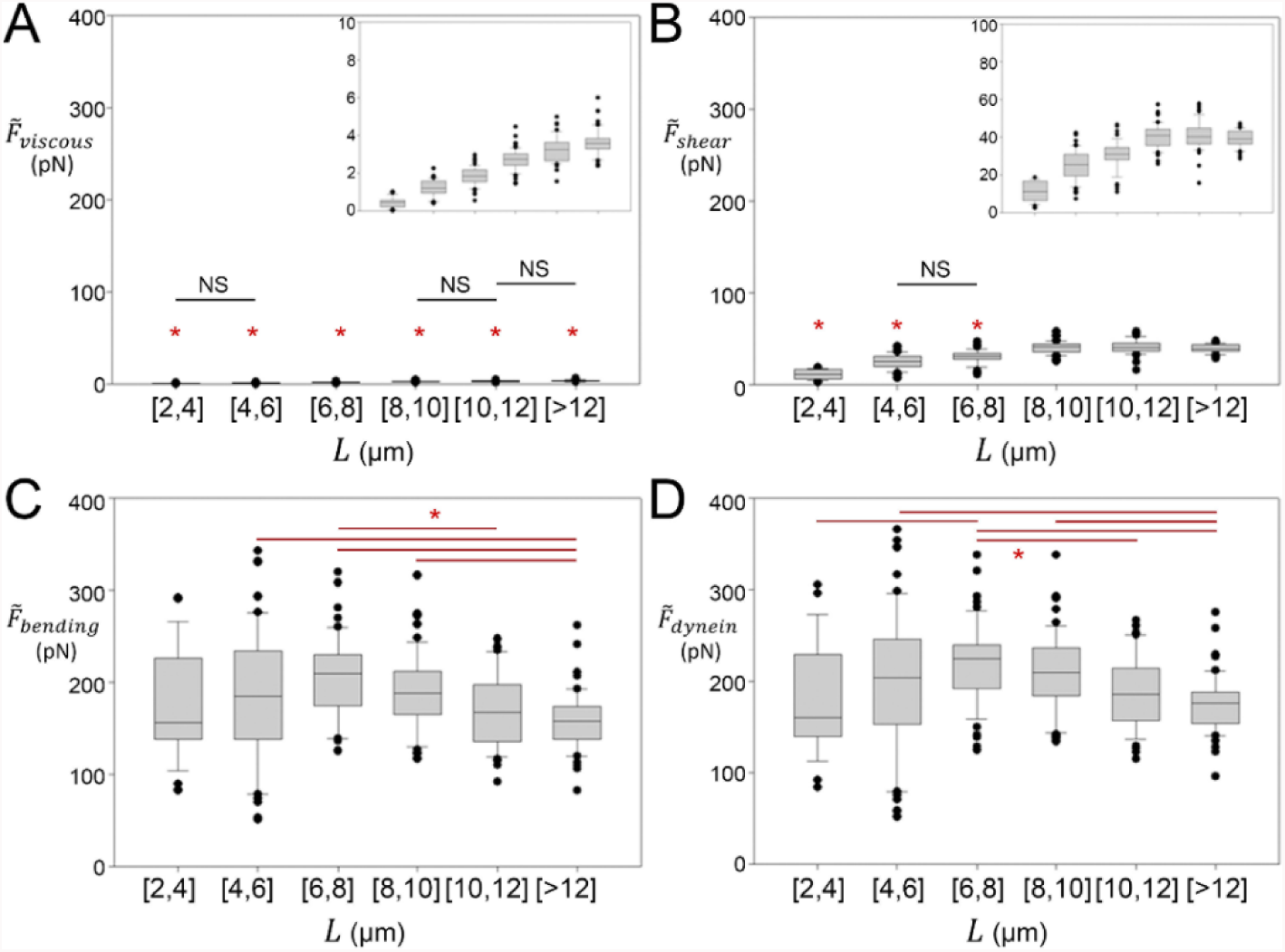
Amplitudes of specific contributions to internal forces within the cilium, estimated from the ciliary waveform, for each length range. (A) Amplitude of force (torque per unit length) required to overcome viscous drag 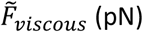. (B) Amplitude of force required to overcome resistance to shear 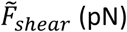. (C) Amplitude of force required to produce elastic bending of doublets and central pair 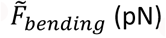. (D) Amplitude of net dynein force 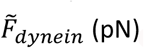.

The values of the dynein force estimated by the current approach (waveform analysis) are comparable to the values for cilia or sperm reported in the literature, as measured by manipulation with glass micro-needles (46–48) or estimated by atomic force microscopy (49, 50). The force per dynein head was estimated from the average net dynein force by invoking several approximations and assumptions. Using the axoneme diameter (*d*_*c*_ = 0.2 *μm*) as the characteristic length associated with the dynein bending moment (48), the dynein force density 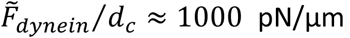 If we assume (conservatively) that only one doublet pairs is active at a time, and that 19 dynein heads (4 triple-headed outer dynein arms, 1 double-headed and 5 single-headed inner dynein arms) repeat every 96 nm, we extrapolate than 198 dynein heads/µm are active. These hypotheses lead to a force of ≈ 5 pN per dynein head. This estimate can be considered an upper bound if multiple doublet pairs are acting synergistically; it is in reasonably good agreement with dynein forces estimated by other methods in mucociliary tissue (50, 51), ciliate outer arms dynein (52), sperm (48, 53, 54) or cytoplasmic dynein (55, 56). Importantly, estimation of the internal force from waveform analysis does not require any type of mechanical perturbation to the cilium.

### Comparison to published mathematical models

To help interpret these measurements in the context of existing hypotheses of waveform generation, cilia motion was simulated using three previously published mathematical models of cilia beating (13, 15, 17) and a simple model of oscillatory motion, a tangentially loaded beam (Supporting Material). Using most parameters from the original publications, none of the mathematical models replicated the experimental data exactly, but each of the models exhibited key features observed in our experimental data (Fig. S3): (i) a critical length was required for periodic beating; (ii) beat frequency was relatively stable (exhibited a plateau) over a range of lengths. While outside the scope of the current experimental study, we expect that each of these models could be refined to better match the critical length and frequency of oscillation, as well as waveform shape. The ability of such models to reproduce trends in behavior as length and other parameters are varied could be addressed systematically in future studies.

### Limitations

#### Uniciliated versus wild-type biciliated *Chlamydomonas*

Uniciliated cells were used to facilitate recording of the waveform, although cilia behavior may differ somewhat in uniciliated compared to biciliated cells. Experiments show that wild-type biciliated *Chlamydomonas* synchronize the two cilia between at least two modes of beating (57–59). Geyer, *et al.* reported that the cilia can be synchronized by “cell-body rocking” with minimal direct hydrodynamic interactions between cilia (60), while others have shown that cilia coordination can be achieved solely through hydrodynamic coupling (61) transmitted to the cell by basal bodies (62). These studies show that ciliary beating in biciliated cells is modulated by factors that are absent in uniciliates. However, cilia of uniciliate cells retain the fundamental ciliary structure and behavior, and in practice are much more convenient for detailed studies of the waveform.

#### Structure and composition of the proximal region

As the cilium regrows, the structure and protein composition of the axoneme might vary. The proximal region of the mature cilium has a different composition than distal regions (63–65). Short cilia during regrowth appear to show some analogous differences (66). While beyond the scope of the current study, a detailed investigation of the ultrastructure and composition of the short, regenerating cilium would complement the current observations.

#### Long-cilia mutant

We did not study cilia behavior in extremely long cilia such as those of *lf* mutants (44, 67, 68). In the current study, using wild-type cells we observed that rotation rate and forces applied by the cilium increased with length. However 10-12 µm cilia had the highest power efficiency ratio, 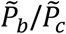 (Fig. 8F); a slight decrease in this ratio was observed for longer cilia (>12 µm). Ciliary beat frequency also decreased slightly for longer (>12 µm) cilia (Fig. 6A). Those results are compatible with the work of Khona, *et al*., on long-cilia mutants (44). In the prior study four *lf* mutants were analyzed and displayed a reduced beat frequency (by 8-16%) as well as a reduced swimming velocity (by 26-57%) for cilia length >18 µm. Taken together, these results suggest that ciliary length is optimized in the normal wild-type cell and that shorter or longer cilia propel the cell with lower efficiency.

## Conclusion

Periodic beating initiates as cilia become longer than 2-4 µm, which suggests that a critical length is necessary for cilia oscillations. In periodically-beating cilia, beat frequency is consistent over the normal range of cilium length, decreasing by a few percent as length increases from 4 µm to 12 µm. Average curvature is also conserved in *Chlamydomonas* cilia at different lengths, so, the waveform shapes of short cilia differ qualitatively from the waveforms of longer cilia. The local mechanical behavior of the cilium is stable over this length range, and thus differences in cilia length alone can explain qualitative changes in the ciliary waveform. In future studies, these data may help distinguish competing hypotheses of the mechanism of ciliary oscillation.

## Author Contributions

MB performed experiments, recorded movies, and analyzed experimental data. KT and PVB developed and solved mathematical models. SKD and PVB designed the research plan. MB, SKD and PVB wrote the manuscript.

## Acknowledgment

We gratefully acknowledge Huawen Lin and Mihaela Stoyanova for their help with deciliation and culture protocols. Support was provided by NSF Grant CMMI-1633971 and the Children’s Discovery Institute of Washington University and Saint Louis Children’s Hospital.

## Supporting Citations

References (69–72) appear in the Supporting Material.

## References

1. Bray, D. 2001. Cell movements : from molecules to motility. Garland Publishing, New York.

2. Wirschell, M., T. Hendrickson, and W. S. Sale. 2007. Keeping an eye on I1: I1 dynein as a model for flagellar dynein assembly and regulation. Cell Motil Cytoskeleton 64(8):569–579.

3. Summers, K. E., and I. R. Gibbons. 1971. Adenosine triphosphate-induced sliding of tubules in trypsin-treated flagella of sea-urchin sperm. Proc Natl Acad Sci U S A 68(12):3092–3096.

4. Satir, P., and T. Matsuoka. 1989. Splitting the ciliary axoneme: implications for a “switch-point” model of dynein arm activity in ciliary motion. Cell Motil Cytoskeleton 14(3):345–358.

5. Nicastro, D., C. Schwartz, J. Pierson, R. Gaudette, M. E. Porter, and J. R. McIntosh. 2006. The molecular architecture of axonemes revealed by cryoelectron tomography. Science 313(5789):944–948.

6. Bui, K. H., T. Yagi, R. Yamamoto, R. Kamiya, and T. Ishikawa. 2012. Polarity and asymmetry in the arrangement of dynein and related structures in the Chlamydomonas axoneme. J Cell Biol 198(5):913–925.

7. Pigino, G., and T. Ishikawa. 2012. Axonemal radial spokes: 3D structure, function and assembly. Bioarchitecture 2(2):50–58.

8. Movassagh, T., K. H. Bui, H. Sakakibara, K. Oiwa, and T. Ishikawa. 2010. Nucleotide-induced global conformational changes of flagellar dynein arms revealed by in situ analysis. Nat Struct Mol Biol 17(6):761–767.

9. Lin, J., T. Heuser, B. I. Carbajal-Gonzalez, K. Song, and D. Nicastro. 2012. The structural heterogeneity of radial spokes in cilia and flagella is conserved. Cytoskeleton (Hoboken) 69(2):88–100.

10. Lin, J., W. Yin, M. C. Smith, K. Song, M. W. Leigh, M. A. Zariwala, M. R. Knowles, L. E. Ostrowski, and D. Nicastro. 2014. Cryo-electron tomography reveals ciliary defects underlying human RSPH1 primary ciliary dyskinesia. Nat Commun 5:5727.

11. Ishikawa, T. 2015. Cryo-electron tomography of motile cilia and flagella. Cilia 4(1):3.

12. Brokaw, C. J. 1972. Flagellar movement: a sliding filament model. Science 178(60):455–462.

13. Hines, M., and J. J. Blum. 1978. Bend propagation in flagella. I. Derivation of equations of motion and their simulation. Biophys J 23(1):41–57.

14. Lin, J., and D. Nicastro. 2018. Asymmetric distribution and spatial switching of dynein activity generates ciliary motility. Science 360(6387).

15. Lindemann, C. B. 1994. A geometric clutch hypothesis to explain oscillations of the axoneme of cilia and flagella. J Theor Biol 168(2):175–189.

16. Riedel-Kruse, I. H., A. Hilfinger, J. Howard, and F. Julicher. 2007. How molecular motors shape the flagellar beat. HFSP J 1(3):192–208.

17. Bayly, P., and S. Dutcher. 2016. Steady dynein forces induce flutter instability and propagating waves in mathematical models of flagella. J Roy Soc Interface 13(123):20160523.

18. Brokaw, C. J. 1972. Computer simulation of flagellar movement. I. Demonstration of stable bend propagation and bend initiation by the sliding filament model. Biophys J 12(5):564–586.

19. Ringo, D. L. 1967. Flagellar motion and fine structure of the flagellar apparatus in Chlamydomonas. J Cell Biol 33(3):543–571.

20. Bayly, P. V., B. L. Lewis, P. S. Kemp, R. B. Pless, and S. K. Dutcher. 2010. Efficient spatiotemporal analysis of the flagellar waveform of Chlamydomonas reinhardtii. Cytoskeleton (Hoboken) 67(1):56–69.

21. Bayly, P. V., B. L. Lewis, E. C. Ranz, R. J. Okamoto, R. B. Pless, and S. K. Dutcher. 2011. Propulsive forces on the flagellum during locomotion of Chlamydomonas reinhardtii. Biophys J 100(11):2716–2725.

22. Tuxhorn, J., T. Daise, and W. L. Dentler. 1998. Regulation of flagellar length in Chlamydomonas. Cell Motil Cytoskeleton 40(2):133–146.

23. Quarmby, L. M. 2004. Cellular deflagellation. International review of cytology 233:47–91.

24. Craige, B., J. M. Brown, and G. B. Witman. 2013. Isolation of Chlamydomonas flagella. Current protocols in cell biology Chapter 3:Unit-3.41.49.

25. Rosenbaum, J. L., J. E. Moulder, and D. L. Ringo. 1969. Flagellar elongation and shortening in Chlamydomonas. The use of cycloheximide and colchicine to study the synthesis and assembly of flagellar proteins. J Cell Biol 41(2):600–619.

26. Yueh, Y. G., and R. C. Crain. 1993. Deflagellation of Chlamydomonas reinhardtii follows a rapid transitory accumulation of inositol 1,4,5-trisphosphate and requires Ca2+ entry. 123(4):869–875.

27. Lefebvre, P. A., S. A. Nordstrom, J. E. Moulder, and J. L. Rosenbaum. 1978. Flagellar elongation and shortening in Chlamydomonas. IV. Effects of flagellar detachment, regeneration, and resorption on the induction of flagellar protein synthesis. J Cell Biol 78(1):8–27.

28. Brokaw, C. J., D. J. Luck, and B. Huang. 1982. Analysis of the movement of Chlamydomonas flagella:” the function of the radial-spoke system is revealed by comparison of wild-type and mutant flagella. J Cell Biol 92(3):722–732.

29. Brokaw, C. J., and D. J. Luck. 1983. Bending patterns of chlamydomonas flagella I. Wild-type bending patterns. Cell Motil 3(2):131–150.

30. Geyer, V. F., P. Sartori, B. M. Friedrich, F. Julicher, and J. Howard. 2016. Independent Control of the Static and Dynamic Components of the Chlamydomonas Flagellar Beat. Curr Biol 26(8):1098–1103.

31. Hyams, J. S., and G. G. Borisy. 1978. Isolated flagellar apparatus of Chlamydomonas: characterization of forward swimming and alteration of waveform and reversal of motion by calcium ions in vitro. 33(1):235–253.

32. Xu, G., K. S. Wilson, R. J. Okamoto, J. Y. Shao, S. K. Dutcher, and P. V. Bayly. 2016. Flexural rigidity and shear stiffness of flagella estimated from induced bends and counterbends. Biophysical Journal 110(12):2759–2768.

33. Holmes, J. A., and S. K. Dutcher. 1989. Cellular asymmetry in Chlamydomonas reinhardtii. J Cell Sci 94 (Pt 2):273–285.

34. Dutcher, S. K. 1995. Flagellar assembly in two hundred and fifty easy-to-follow steps. Trends Genet 11(10):398–404.

35. Lux, F. G., 3rd, and S. K. Dutcher. 1991. Genetic interactions at the FLA10 locus: suppressors and synthetic phenotypes that affect the cell cycle and flagellar function in Chlamydomonas reinhardtii. Genetics 128(3):549–561.

36. Brokaw, C. J., and D. J. Luck. 1985. Bending patterns of chlamydomonas flagella: III. A radial spoke head deficient mutant and a central pair deficient mutant. Cell Motil 5(3):195–208.

37. Schneider, C. A., W. S. Rasband, and K. W. Eliceiri. 2012. NIH Image to ImageJ: 25 years of image analysis. Nature methods 9(7):671–675.

38. Gray, J., and G. J. Hancock. 1955. The Propulsion of Sea-Urchin Spermatozoa. 32(4):802–814.

39. Oberbeck, A. 1876. Ueber stationäre Flüssigkeitsbewegungen mit Berücksichtigung der inneren Reibung. Journal für die reine und angewandte Mathematik 81:62–80.

40. Chwang, A. T., and T. Y.-T. Wu. 1975. Hydromechanics of low-Reynolds-number flow. Part 2. Singularity method for Stokes flows. Journal of Fluid Mechanics 67(4):787–815.

41. Minoura, I., T. Yagi, and R. Kamiya. 1999. Direct Measurement of Inter-doublet Elasticity in Flagellar Axonemes. Cell Structure and Function 24(1):27–33.

42. Avasthi, P., M. Onishi, J. Karpiak, R. Yamamoto, L. Mackinder, Martin C. Jonikas, Winfield S. Sale, B. Shoichet, John R. Pringle, and Wallace F. Marshall. 2014. Actin Is Required for IFT Regulation in Chlamydomonas reinhardtii. Current Biology 24(17):2025–2032.

43. Hunter, E. L., W. S. Sale, and L. M. Alford. 2016. Analysis of Axonemal Assembly During Ciliary Regeneration in Chlamydomonas. Methods in molecular biology (Clifton, N.J.) 1454:237–243.

44. Khona, D. K., V. G. Rao, M. J. Motiwalla, P. C. Varma, A. R. Kashyap, K. Das, S. M. Shirolikar, L. Borde, J. A. Dharmadhikari, A. K. Dharmadhikari, S. Mukhopadhyay, D. Mathur, and J. S. D’Souza. 2013. Anomalies in the motion dynamics of long-flagella mutants of Chlamydomonas reinhardtii. Journal of biological physics 39(1):1–14.

45. Oltean, A., A. J. Schaffer, P. V. Bayly, and S. L. Brody. 2018. Quantifying Ciliary Dynamics during Assembly Reveals Stepwise Waveform Maturation in Airway Cells. Am J Respir Cell Mol Biol 59(4):511–522.

46. Yoneda, M. 1960. Force Exerted by a Single Cilium of Mytilus Edulis. I. 37(3):461–468.

47. Tani, T., and S. Kamimura. 1999. Dynein-ADP as a force-generating intermediate revealed by a rapid reactivation of flagellar axoneme. Biophysical journal 77(3):1518–1527.

48. Schmitz, K. A., D. L. Holcomb-Wygle, D. J. Oberski, and C. B. Lindemann. 2000. Measurement of the force produced by an intact bull sperm flagellum in isometric arrest and estimation of the dynein stall force. Biophysical journal 79(1):468–478.

49. Sakakibara, H. M., Y. Kunioka, T. Yamada, and S. Kamimura. 2004. Diameter oscillation of axonemes in sea-urchin sperm flagella. Biophys J 86(1 Pt 1):346–352.

50. Teff, Z., Z. Priel, and L. A. Gheber. 2007. Forces applied by cilia measured on explants from mucociliary tissue. Biophysical journal 92(5):1813–1823.

51. Hill, D. B., V. Swaminathan, A. Estes, J. Cribb, E. T. O’Brien, C. W. Davis, and R. Superfine. 2010. Force generation and dynamics of individual cilia under external loading. Biophys J 98(1):57–66.

52. Hirakawa, E., H. Higuchi, and Y. Y. Toyoshima. 2000. Processive movement of single 22S dynein molecules occurs only at low ATP concentrations. Proc Natl Acad Sci U S A 97(6):2533–2537.

53. Kamimura, S., and K. Takahashi. 1981. Direct measurement of the force of microtubule sliding in flagella. Nature 293:566.

54. Shingyoji, C., H. Higuchi, M. Yoshimura, E. Katayama, and T. Yanagida. 1998. Dynein arms are oscillating force generators. Nature 393(6686):711–714.

55. Ashkin, A., K. Schutze, J. M. Dziedzic, U. Euteneuer, and M. Schliwa. 1990. Force generation of organelle transport measured in vivo by an infrared laser trap. Nature 348(6299):346–348.

56. Toba, S., T. M. Watanabe, L. Yamaguchi-Okimoto, Y. Y. Toyoshima, and H. Higuchi. 2006. Overlapping hand-over-hand mechanism of single molecular motility of cytoplasmic dynein. 103(15):5741–5745.

57. Rüffer, U., and W. Nultsch. 1987. Comparison of the beating of cis- and trans-flagella of Chlamydomonas cells held on micropipettes. Cell Motility 7(1):87–93.

58. Polin, M., I. Tuval, K. Drescher, J. P. Gollub, and R. E. Goldstein. 2009. Chlamydomonas swims with two “gears” in a eukaryotic version of run-and-tumble locomotion. Science 325(5939):487–490.

59. Leptos, K. C., K. Y. Wan, M. Polin, I. Tuval, A. I. Pesci, and R. E. Goldstein. 2013. Antiphase Synchronization in a Flagellar-Dominance Mutant of Chlamydomonas. Physical Review Letters 111(15):158101.

60. Geyer, V. F., F. Jülicher, J. Howard, and B. M. Friedrich. 2013. Cell-body rocking is a dominant mechanism for flagellar synchronization in a swimming alga. 110(45):18058–18063.

61. Brumley, D. R., K. Y. Wan, M. Polin, and R. E. Goldstein. 2014. Flagellar synchronization through direct hydrodynamic interactions. eLife 3:e02750.

62. Wan, K. Y., and R. E. Goldstein. 2016. Coordinated beating of algal flagella is mediated by basal coupling. 113(20):E2784–E2793.

63. Piperno, G., and Z. Ramanis. 1991. The proximal portion of Chlamydomonas flagella contains a distinct set of inner dynein arms. 112(4):701–709.

64. Bui, K. H., H. Sakakibara, T. Movassagh, K. Oiwa, and T. Ishikawa. 2009. Asymmetry of inner dynein arms and inter-doublet links in Chlamydomonas flagella. The Journal of cell biology 186(3):437–446.

65. Yagi, T., K. Uematsu, Z. Liu, and R. Kamiya. 2009. Identification of dyneins that localize exclusively to the proximal portion of Chlamydomonas flagella. 122(9):1306–1314.

66. Satish Tammana, T. V., D. Tammana, D. R. Diener, and J. Rosenbaum. 2013. Centrosomal protein CEP104 (Chlamydomonas FAP256) moves to the ciliary tip during ciliary assembly. 126(21):5018–5029.

67. Barsel, S. E., D. E. Wexler, and P. A. Lefebvre. 1988. Genetic analysis of long-flagella mutants of Chlamydomonas reinhardtii. Genetics 118(4):637–648.

68. Asleson, C. M., and P. A. Lefebvre. 1998. Genetic analysis of flagellar length control in Chlamydomonas reinhardtii: a new long-flagella locus and extragenic suppressor mutations. Genetics 148(2):693–702.

69. Bayly, P. V., and K. S. Wilson. 2014. Equations of inter-doublet separation during flagella motion reveal mechanisms of wave propagation and instability Biophys J 107(7):1756–1772.

70. Bolotin, V. V. 1963. Nonconservative problems of the theory of elastic stability. Macmillan, New York,.

71. Leipholz, H. H. E. 1987. Stability theory : an introduction to the stability of dynamic systems and rigid bodies. B.G. Teubner - Wiley, Stuttgart - Chichester ; New York.

72. Virgin, L. N. 2007. Vibration of Axially-Loaded Structures. Cambridge University Press, New York.

